# A platform for semi-automated voluntary training of common marmosets for behavioral neuroscience: Voluntary training of common marmosets

**DOI:** 10.1101/635334

**Authors:** Jeffrey D. Walker, Friederice Pirschel, Nicholas Gidmark, Jason N. MacLean, Nicholas G. Hatsopoulos

## Abstract

In most cases, behavioral neuroscience studies of the common marmoset employ adaptations of well-established methods used with macaque monkeys. However, in most cases these approaches do not readily generalize to marmosets indicating a need for alternatives. Here we present the development of one such alternate: a platform for semi-automated, voluntary in-home cage behavioral training that allows for the study of naturalistic behaviors. We describe the design and production of a modular behavioral training apparatus using CAD software and digital fabrication. We demonstrate that this apparatus permits voluntary behavioral training and data collection throughout the marmoset’s waking hours with little experimenter intervention. Further we demonstrate the use of this apparatus to reconstruct the kinematics of the marmoset’s upper limb movement during natural foraging behavior.

**NEW AND NOTEWORTHY:** The study of marmosets in neuroscience has grown rapidly and this model organism presents challenges that are unique to this primate species. Here we address those challenges with an innovative platform for semi-automated and voluntary training of common marmosets. The platform allows marmosets to train throughout their waking hours with little to no experimenter intervention. We describe the use of this platform to capture the kinematics of the upper limb during natural foraging behavior and to expand the opportunities for behavioral training beyond the limits of traditional behavioral training sessions. The platform is flexible and can be easily extended to incorporate other motor tasks (e.g. visually cued reaching or manipulandum based tasks) using CAD models and digital fabrication.

## INTRODUCTION

Neurophysiological recordings of isolated single neurons in awake, behaving macaques began in the late 1960’s, whereas the first reports of single neuron recordings from awake marmosets did not occur until the early 2000s (Evarts, 1968; Lu et al., 2001). Despite the relative recency of broad adoption of the marmoset as a model species for systems neuroscience there is growing interest, but the techniques for working with marmosets in this context are relatively new as compared to those used with more standard model primate species (e.g. rhesus macaques) in neuroscience research. Because of the success of the model, the approach to training a macaque to perform an experimental task has remained, with few exceptions, relatively unchanged for decades. In general, the monkey is restrained while engaging in a trained task for a few hours in exchange for water or juice. This method is popular because it generally yields hundreds to thousands of repetitions of a given behavior over the course of a training session. However, our experience and the early behavioral work indicate that this approach may be ill-suited for working with marmosets. It yields far fewer trials and limits the expression of natural behavior (Johnston et al. 2017; Prins et al., 2017; Eliades and Wang, 2003). To partly address these issues, Wang and colleagues developed a technique for wireless neural recordings which allowed for the study of sensorimotor processing in freely vocalizing marmosets (Roy and Wang, 2012). However, there has not been a complimentary innovation in behavioral training paradigms to increase trial counts.

### Marmoset ethology and its implications for experimental design

Marmosets are obligate gum feeders and prey species. Field studies estimate that marmosets spend about 30 percent of their waking hours feeding on exudates (Maier et al., 1982 in Sussman and Kinzey 1984) and spend 25-30% of their waking time foraging for insects (Abreu et al., 2016; Stevenson and Rylands, 1988). In order to feed on exudates, marmosets must gouge wounds into the trunks of trees to access the gum. They gouge new holes and revisit previously gouged holes to feed on newly accumulated gum (Stevenson and Rylands, 1988). These visits only last a few seconds (Stevenson and Rylands, 1988). Their daily behavioral repertoire generally does not involve them sitting in a single place engaging in repetitive behaviors for multiple hours. With this in mind, we designed an approach to training marmosets that would allow them to voluntarily engage in experimental behavior for short sessions throughout their waking hours. In order to do so we sought to modify an approach successful applied to rodents where rats voluntarily head-fixed themselves for in vivo calcium imaging (Scott, Brody, and Tank 2013). To implement this approach, researchers designed a set of custom elements to ensure stable imaging and slowly acclimated the rat to the apparatus, gradually extending the duration of head fixation. Once the animal was trained, the process of data collection could proceed with minimal experimenter involvement. This sort of voluntary setup, that allowed the animal to engage in the experiment throughout the day as an expression of its normal behavioral repertoire, seemed like a promising approach to behavioral training of marmosets.

## MATERIALS AND METHODS

### Subjects

All work described were done with three common marmosets (Callithrix jacchus) (two females, and one male, 375-410 g). All methods were approved by the Institutional Animal Care and Use Committee of the University of Chicago.

### Design criteria

Informed by field studies of the marmoset’s natural behavioral repertoire (Stevenson and Rylands, 1988; Sussman and Kinzey, 1984), early work with marmosets in neuroscience (Eliades and Wang, 2003, 2005, 2008a), and the novel approach to training and in-vivo calcium imaging developed by Scott, Brody, and Tank (2013), we developed a behavioral training apparatus that attaches to the marmosets’ home cage. This apparatus allows marmosets to voluntarily engage in behavioral training throughout their waking hours.

The three primary design criteria for the final apparatus were 1) that it mounts to the home cage to allow for voluntary engagement in training throughout the marmosets’ waking hours, 2) that it provides reliable positioning of marmosets and clear views of the upper limbs for capturing the kinematics of reaching movements, and 3) it provides a flexible way to present different experimental tasks. Additionally, to validate the effectiveness of the apparatus as a training instrument, it had to have a way to monitor and record the marmosets’ behavior within it. Finally, to facilitate training using operant conditioning, the apparatus also had to include a method for precisely timed reward delivery.

### Hardware design and iteration

Inspired by the gum feeding behavior in which marmosets naturally engage, the first version of the apparatus trained the marmosets to assume the appropriate posture to receive a small volume of yogurt (Figure 1A). This posture placed them in front of a tray that contained foraging substrate. The next version of the apparatus removed the yogurt reward, and we found that marmosets would still engage in foraging behavior within the apparatus. After a series of iterations optimizing the form of the apparatus to multiple motion capture modalities, the current version of the apparatus (Figure 1B) has allowed us to record the kinematics of this foraging behavior to study sensorimotor cortical responses related to upper limb movement.

**Figure 1.**
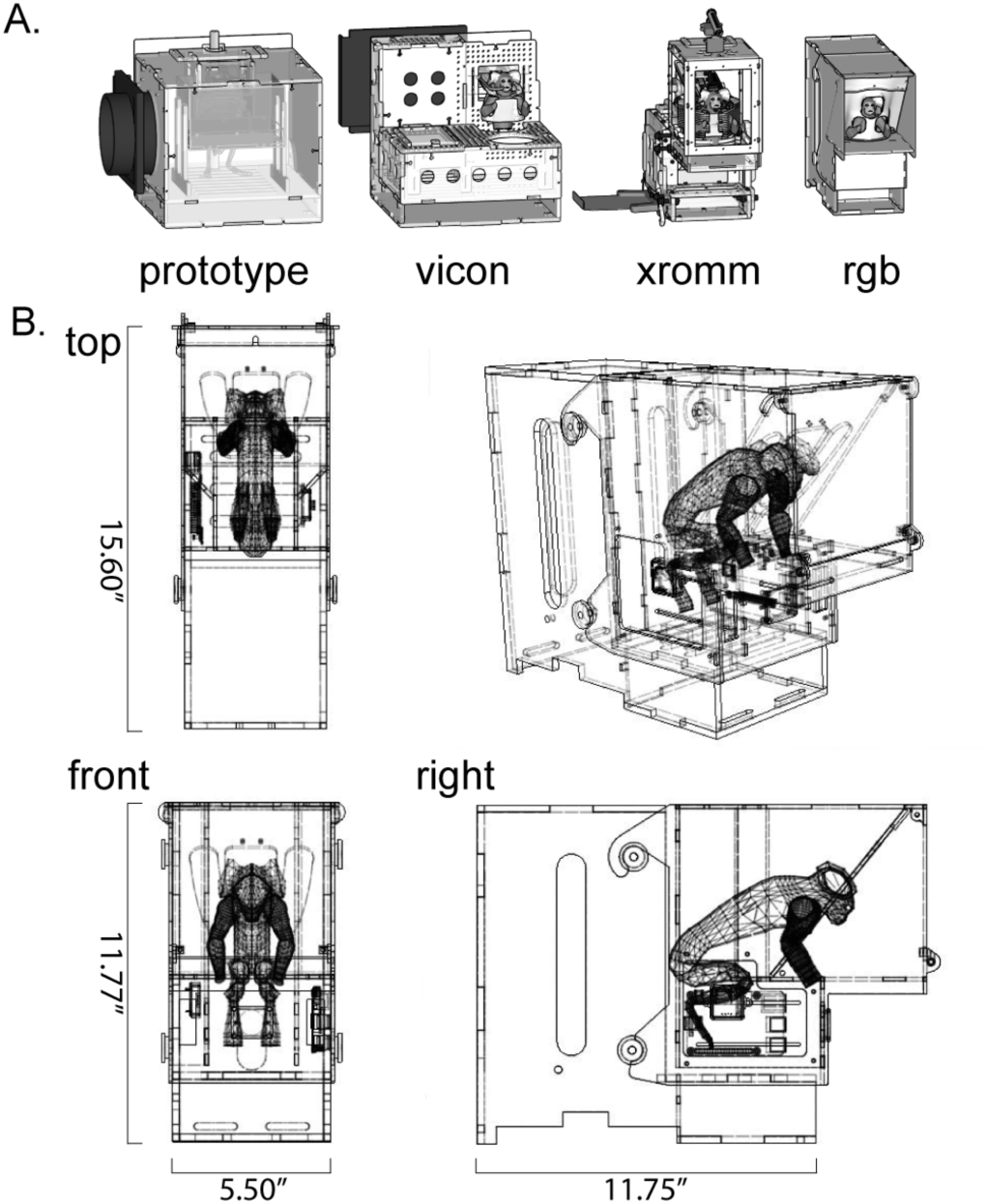
Developing a voluntary in-home cage approach to behavioral training with marmosets. A) Iterations of apparatus design optimized for different motion capture modalities. B) Drawing of current version of the behavioral training apparatus.

Designs of early versions of the behavioral training apparatus were done with 3D CAD software called SketchUp, while later versions were designed using AutoDesk Fusion 360 (Figure 1). The core of the apparatus was constructed using 1/8” or 1/4” thick clear acrylic sheets (continuous cast, McMaster Carr, Elmhurst, IL) that were cut into interlocking panels using a laser cutter (Universal Laser Systems VLS4.60). These panels were then assembled to achieve the form of the apparatus. To monitor the activity of the marmosets within the apparatus, we designed a simple circuit (Figure 2A-B) that included two photocells (CdS - photoresistor) and one infrared light based switch (IR switch comprising an IR phototransistor and IR LED pair), a syringe pump (syringepump.com, NE-500) and a network-connected microcontroller (Arduino YÚN). The sensors acted as triggers to log the marmosets entering and leaving the gate, the belt, and the nosepiece of the apparatus. The sensor readings were logged to an SD card within the microcontroller, and the network connection of the microcontroller allowed remote operation of the apparatus. The apparatus sat on top of a round gate installed in ceiling of the home cage (part # 1822K314, McMaster-Carr).

**Figure 2.**
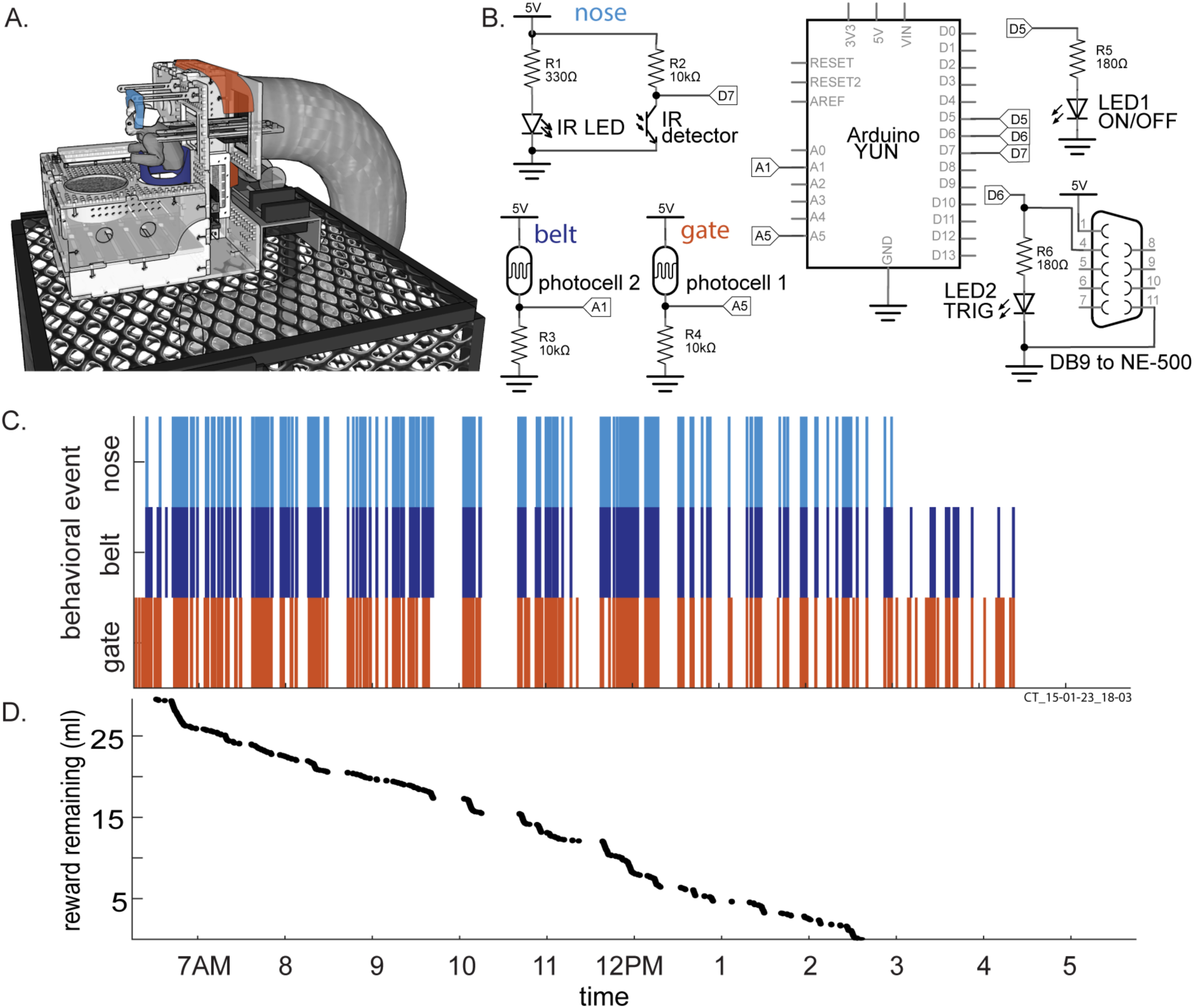
Hardware design and single day of behavior within the apparatus. A) Illustration of behavioral training apparatus with sensors embedded into the gate (orange), belt (dark blue) and nosepiece (light blue) to log behavior throughout the day. The foraging tray is placed in front of the belt. A syringe pump is connected to deliver reward. The whole assembly sits on top of the home cage. B) Circuit diagram detailing the circuit logging behavior and delivering reward. C and D) Results for single day of behavior within the apparatus. C) Vertical ticks indicate the time of trigger events for sensors within the gate, belt and nosepiece. For instance, an orange tick indicates the marmoset crossed the gate of the apparatus, a dark blue tick indicates the marmoset is within the belt of the apparatus and a light blue tick indicates that the marmoset has its nose positioned within the nosepiece. When the marmoset stays within the nosepiece, 0.1 ml of yogurt is dispensed every 10 seconds as positive reinforcement for assuming the appropriate posture. D) Reward remaining as a function of time of day.

### Software for automating and monitoring training

We wrote a library (C++) to coordinate logging activity within the apparatus, evaluate reward-conditions, deliver reward, and allow remote apparatus operation. The object-oriented design of this library is meant to facilitate integration of future experimental tasks.

## RESULTS

### Foraging

We began by studying foraging since this was a behavior in which the marmosets readily engaged. Foraging was coupled with the task of assuming an appropriate posture in exchange for yogurt reward. Marmosets engaged in behavior within the training apparatus throughout the day, and their engagement was sensitive to reward availability (Figure 2C-D). We measured each time a marmoset entered and exited the belt of the apparatus, i.e. the start and end of a session, to quantify the duration of these sessions. This measure allowed us to generate an estimate of how much time marmosets would spend engaging in behavior within the apparatus and how that behavior was distributed throughout their waking hours. Over the course multiple days, we found that marmosets would spend up to an hour each day engaging in behavior within the apparatus spanning 65 – 216 sessions (Figure 3A-B). Most of these sessions were not longer than 20 seconds, but some lasted almost five minutes (n= 1739 sessions over 11 days, mean = 17.00 sec, median = 9.36 sec, Q1 = 6.99 sec, Q3 = 16.67, max = 270 sec) (Figure 3C-D).

**Figure 3.**
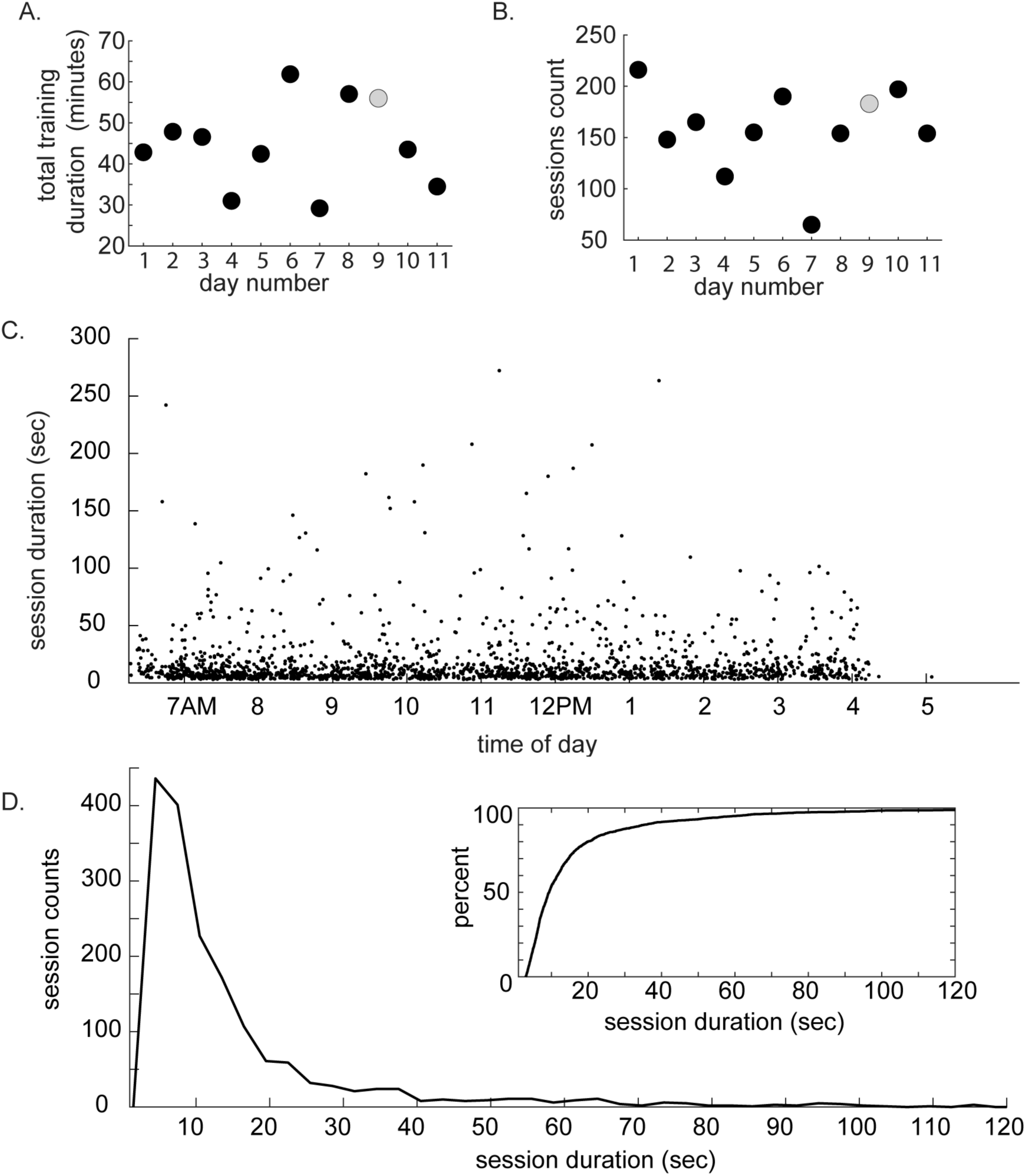
Summary of sessions of behavior within the apparatus across days. A) Total duration of all sessions within each day. B) Number of sessions of behavior within each day. Grey circle indicates data point corresponding to day illustrated in Figure 2. C) Session durations as a function of time of day. Data were pooled across all days. Each point represents a single session. D) Distribution of session durations. Inset: cumulative distribution of session durations.

After validating that marmosets engage in voluntary behavioral training and gaining a sense of their attention span, we optimized the design of the apparatus to provide unobscured views of the upper limb (Figure 1) and set out to characterize foraging behavior within the apparatus. Using a custom-written algorithm (MATLAB) to define the video frame when the animal started foraging, reaches were subsequently counted manually. Reaches with both hands were counted over the course of a day (12 hours). When only the foraging mix was provided in the apparatus, marmosets performed between 20 and 80 reaches while foraging each day (Figure 4A). In contrast, if their entire daily diet (i.e. foraging mix and normal diet) was provided within the behavioral training apparatus, the marmosets performed 100-300 reaches per day (Figure 4B).

**Figure 4.**
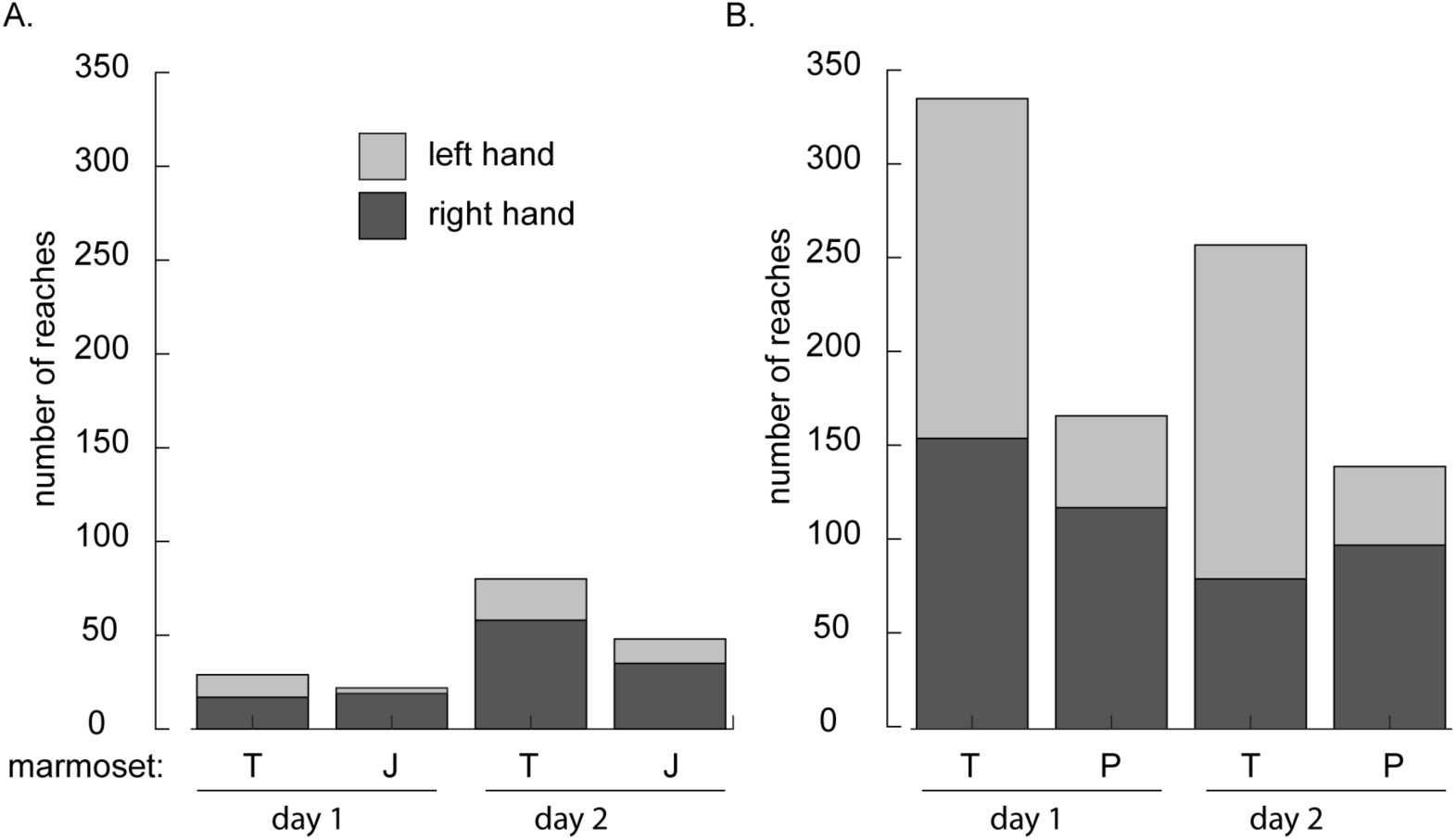
Reaches counted during foraging behavior within the apparatus. A) Counts of reaches for two marmosets (T and J) across two days when only foraging mix was provided in the apparatus. B) Counts of reaches for two marmosets (T and P) across two days when their entire daily diet was provided within the training apparatus.

### Recording upper limb kinematics during foraging with XROMM

We next sought to record the kinematics of the upper limb during foraging. After confirming that marmosets do not tolerate retro-reflective markers placed on their skin needed for traditional near infrared based motion capture systems (e.g. VICON) (Takemi et al., 2014; Young et al., 2016) we moved to using an x-ray based system called XROMM, or X-Ray Reconstruction of Moving Morphology (Brainerd et al. 2010) (Figure 5A). Bi-planar x-ray sources and image intensifiers (90 kV, 25 mA at 200fps) allowed us to reconstruct time varying joint angles by tracking the 3D position of radio-opaque tantalum beads (0.5-1 mm, Bal-tec) placed within the soft tissue of the arm, hand and torso (Figure 5B). Using a set of tools developed at Brown University (Brainerd et al., 2010; Knörlein et al., 2016; Miranda et al., 2011), and adaptations of joint coordinate systems for the upper limb (Baier and Gatesy, 2013; Wu et al., 2005), we could translate the position of these markers into joint kinematics (Figure 5). We placed markers in the torso and upper limb subcutaneously using angiocatheters (16G, Becton, Dickinson and Company) (Figure 5B). The marker set illustrated allowed reconstruction of the seven degrees of freedom of the shoulder, elbow and wrist (Figure 5C).

**Figure 5.**
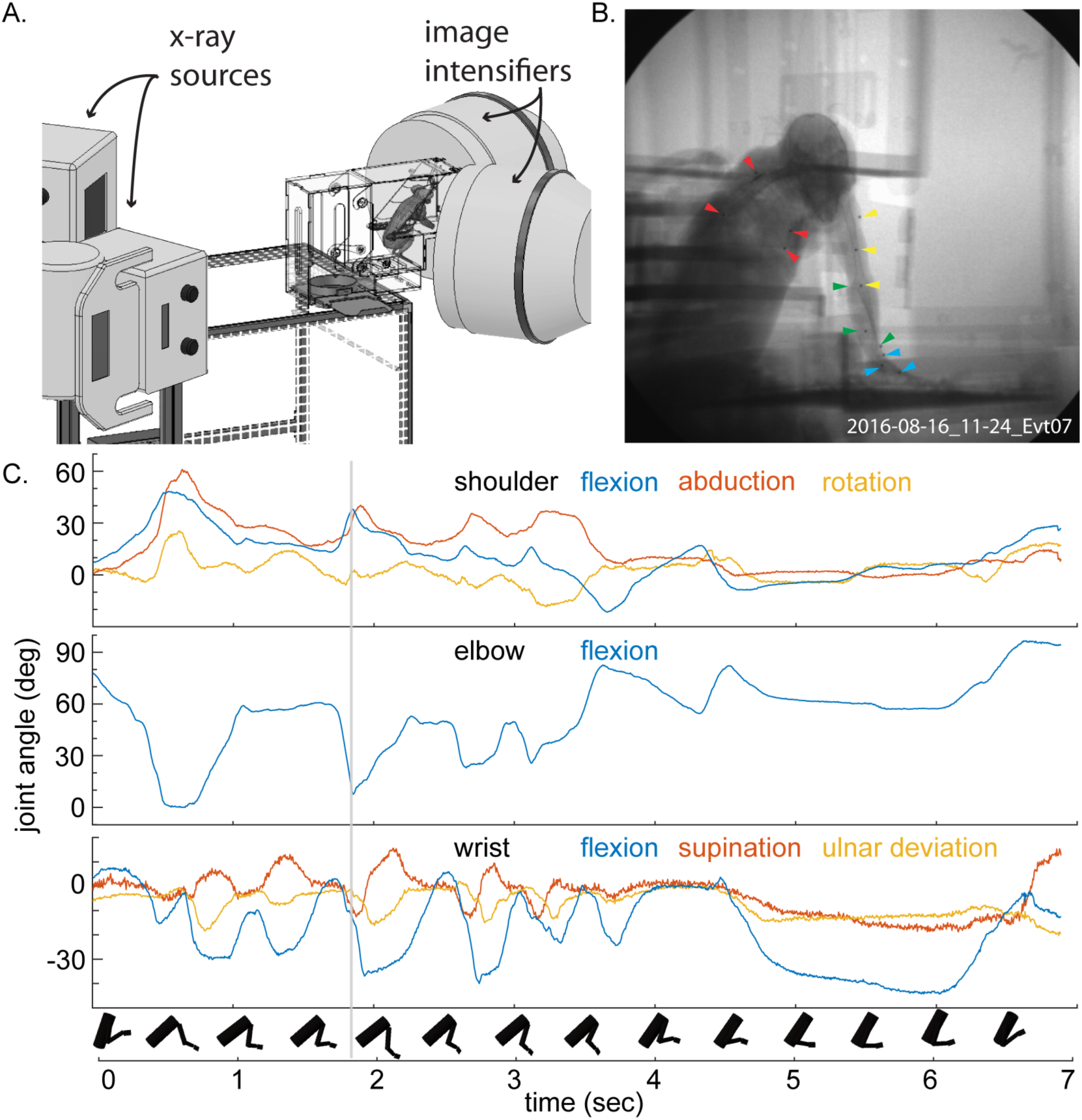
Capturing the kinematics of upper limb during foraging with XROMM. A) An illustration of the XROMM bi-planar x-ray motion capture system together with the behavioral training apparatus and marmoset in the capture volume. B) A single frame of x-ray video of a marmoset foraging within the apparatus. Note radio-opaque markers placed within the marmoset’s torso (red), upper arm (yellow), forearm (green) and hand (blue). C) Seven degrees of freedom of upper limb movement reconstructed by tracking the movement of the radio-opaque markers seexsn in B). Grey line indicates timestamp of the frame in B). Rigid bodies represent kinematics of the torso, upper arm, forearm and hand over the course of the foraging sequence.

## DISCUSSION

Here we present a method for in home-cage, semi-automated, and voluntary behavioral training of marmosets that has in our experience been more prolific than adaptations of the more traditional approaches. The method presented also allows for the training of multiple marmosets in parallel. It provides a flexible platform for a variety of experimental tasks and liberates the animals from excessive restraints and provides a platform for marmosets to self-initiate natural behavior in addition to engaging in more traditional operant paradigms. It allows for behavioral engagement in short sessions throughout the marmosets waking hours rather than extended sessions, which are limited by marmoset cooperation and satiation. This flexible approach should allow us to contextualize results from constrained and over-trained experimental tasks within the space of the marmoset’s natural behavioral repertoire. Toward this end, we are in the process of implementing an additional motor learning task and we are pairing this training approach with wireless neural recordings.

It is clear that marmosets are well poised to contribute to our understanding of the operating principles of neocortex as attested by their increasing prevalence in published systems neuroscience reports (Miller, 2017). Moreover the structure of marmoset neocortex provides a strong potential for targeted circuit manipulations (Belmonte et al., 2015; Sasaki et al., 2009). Marmosets have been trained to perform experimental tasks such as eye fixation and a smooth pursuit (Mitchell, Reynolds, and Miller 2014; Mitchell, Priebe, and Miller 2015) and basic reaching and neuroprosthetic tasks (Ebina et al., 2018; Pohlmeyer et al. 2012, 2014) using training procedures common in macaque studies. But the quantity of behavior marmosets produce using these procedures is generally limited in comparison to that of macaques. In contrast, the techniques we designed dramatically increased the time available for behavioral training by eliminating the use of restraint and making the experimental training apparatus available to the marmosets throughout their waking hours. With this paradigm, our initial estimates suggest that we can minimally double and can often quadruple the quantity of experimentally useful behavioral trials with the added benefit that this behavior is self-initiated rather than generated through restriction. We would like to see if, with adaptations such as support for reliable eye positioning, this behavioral training approach could be useful to increase the behavioral output of marmosets in studies of other systems. Finally, as we have argued (Walker et al., 2017), natural behaviors in of themselves warrant study and this particular experimental behavioral paradigm facilitates this class of study.

## CODE AND DESIGN FILE ACCESSIBILITY

Both software written to coordinate training and design files used to fabricate the apparatus are available from the authors upon request.

## Acknowledgements

We thank Marek Niekrasz and the veterinary staff at the University of Chicago for their assistance with marmoset care and Callum Ross for assistance with the XROMM system. This work was supported by NIH R01 NS104898, NSF MRI 1338066, NSF IGERT, UChicago Big Vision Fund and the Tarrson Fund.

